# Latent variable model for aligning barcoded short-reads improves downstream analyses

**DOI:** 10.1101/220236

**Authors:** Ariya Shajii, Ibrahim Numanagić, Bonnie Berger

**Affiliations:** Computer Science and AI Lab, Massachusetts Institute of Technology, Cambridge, MA, USA; Department of Mathematics, Massachusetts Institute of Technology, Cambridge, MA, USA

## Abstract

Recent years have seen the emergence of several “third-generation” sequencing platforms, each of which aims to address shortcomings of standard next-generation short-read sequencing by producing data that capture long-range information, thereby allowing us to access regions of the genome that are inaccessible with short-reads alone. These technologies either produce physically longer reads typically with higher error rates or instead capture long-range information at low error rates by virtue of read “barcodes” as in 10x Genomics’ Chromium platform. As with virtually all sequencing data, sequence alignment for third-generation sequencing data is the foundation on which all downstream analyses are based. Here we introduce a latent variable model for improving barcoded read alignment, thereby enabling improved downstream genotyping and phasing. We demonstrate the feasibility of this approach through developing EMerAld— or EMA for short— and testing it on the barcoded short-reads produced by 10x’s sequencing technologies. EMA not only produces more accurate alignments, but unlike other methods also assigns interpretable probabilities to the alignments it generates. We show that genotypes called from EMA’s alignments contain over 30% fewer false positives than those called from Lariat’s (the current 10x alignment tool), with a fewer number of false negatives, on datasets of NA12878 and NA24385 as compared to NIST GIAB gold standard variant calls. Moreover, we demonstrate that EMA is able to effectively resolve alignments in regions containing nearby homologous elements— a particularly challenging problem in read mapping— through the introduction of a novel statistical binning optimization framework, which allows us to find variants in the pharmacogenomically important *CYP2D* region that go undetected when using Lariat or BWA. Lastly, we show that EMA’s alignments improve phasing performance compared to Lariat’s in both NA12878 and NA24385, producing fewer switch/mismatch errors and larger phase blocks on average.

EMA software and datasets used are available at http://ema.csail.mit.edu.

## 1 Introduction

As sequencing technologies have continued to advance beyond the initial introduction of next-generation sequencing (NGS), we have begun to see the emergence of so-called “third-generation” sequencing platforms, which seek to improve on the standard short-read sequencing that has thus far been the heart of most next-generation sequencing technologies [1]. Several organizations are at the center of this new sequencing revolution, including Pacific Biosciences [2], Oxford Nanopore [3] and 10x Genomics [4]. While the former two have developed sequencing methods that produce much longer physical reads (e.g., 10–200kb) at typically higher error rates, 10x Genomics’ Chromium platform instead generates reads that have long-range information implicit in a read “barcode,” a 16bp nucleic acid sequence that can be used to determine the source fragment of a particular read [5]. At a high level, 10x sequencing first organizes long (10–200kb) DNA fragments into droplets such that few are present in each droplet. These fragments are then sheared, and a unique barcoded bead is added to each droplet so as to ligate a 16bp barcode to each sheared piece. These newly-barcoded sheared fragments are then sequenced using standard short-read sequencing, thereby producing barcoded short-reads [4]. Because they help identify the original source fragment, these barcodes implicitly carry long-range information, and can have a significant impact on alignment as well as fundamental downstream analyses such as structural variation detection and phasing.

Barcoded reads have several advantages over physically long reads. Firstly, and perhaps most importantly, barcoded short-read sequencing is substantially cheaper than long-read sequencing; whereas PacBio’s and Oxford Nanopore’s sequencing platforms currently cost anywhere from $750–$1000 per GB of data, 10x sequencing is a comparatively cheap add-on to standard short-read sequencing, and therefore bears the same cost (specifically, $30 per GB) plus a $500 overhead per sample [5]. Secondly, the error profile of barcoded short-reads is very similar to those of standard short-reads (roughly 0.1% substitution errors), which enables us to augment the tools and algorithms that have been developed for regular short-reads to handle their barcoded counterparts. By contrast, long-read sequencing typically produces high rates of erroneous indels (ranging from 12–13%), which presents a challenge when trying to use preexisting algorithms.

As with virtually all sequencing data, the first step in the analysis pipeline for barcoded reads is typically alignment. While barcoded reads can in theory be aligned by a standard short-read aligner (e.g, CORA [6], BWA [7], Bowtie2 [8]), this would fail to take advantage of the information provided by the barcodes. An alternative approach given by McCoy *et al.* is to assemble the reads for each particular barcode, and to treat the result as a single “synthetic long-read” [9]. While this works well for technologies like Illumina’s TruSeq Synthetic Long-Read platform (formerly Moleculo)— which is similar to 10x sequencing except that source fragments are sequenced with a much higher coverage— the fact that 10x sequencing achieves its coverage not by having high per-droplet coverage, but rather by having many droplets, makes such assembly impractical for 10x data. On the other hand, the fact that TruSeq sequences fragments at high coverage inflates their sequencing costs to be on par with PacBio’s and Oxford Nanopore’s, whereas 10x circumvents this high cost via shallow fragment sequencing [5].

Currently, the state-of-the-art in terms of barcoded read alignment employs “read clouds”— groups of reads that share the same barcode and map to the same genomic region— to choose the most likely alignment from a set of candidate alignments for each read [10]. Intuitively, read clouds represent the possible source fragments from which the barcoded reads are derived. The read cloud approach to alignment effectively begins with a standard all-mapping to a reference genome to identify these clouds, followed by an iterative update of reads’ assignments to their possible alignments, guided by a Markov random field that is used to evaluate the probability of a given configuration, taking into account the alignment scores, clouds, etc. Moreover, with 10x technology, as multiple fragments can share the same barcode, it is in general not possible to infer the source fragment for a read (and thus its correct alignment within a reference genome) merely by looking at its barcode. In order to deduce the correct placement of a read (and thus its unknown source fragment), all possible alignments of that read (and thus its possible fragments) need to be examined.

Here, we propose a new probabilistic optimization framework for barcoded read alignment that employs a probabilistic interpretation of clouds: EMerAld, or EMA for short (Figure 1). Our two main conceptual advances are as follows. Intuitively, rather than assigning each read to just one of its possible alignments at any given time, we make use of probabilistic assignments and employ a latent variable model to determine final alignment probabilities and, ultimately, to select the most likely alignments. During the alignment process, we utilize a disjoint-set data structure over read clouds to normalize alignment probabilities in a physically sensible way. Secondly, we propose a statistical binning optimization approach to better handle the ubiquitous repetitive regions of the genome, compared to the currently-used method of simply picking the lowest edit distance alignment of a read in a given cloud.

**Figure 1:**
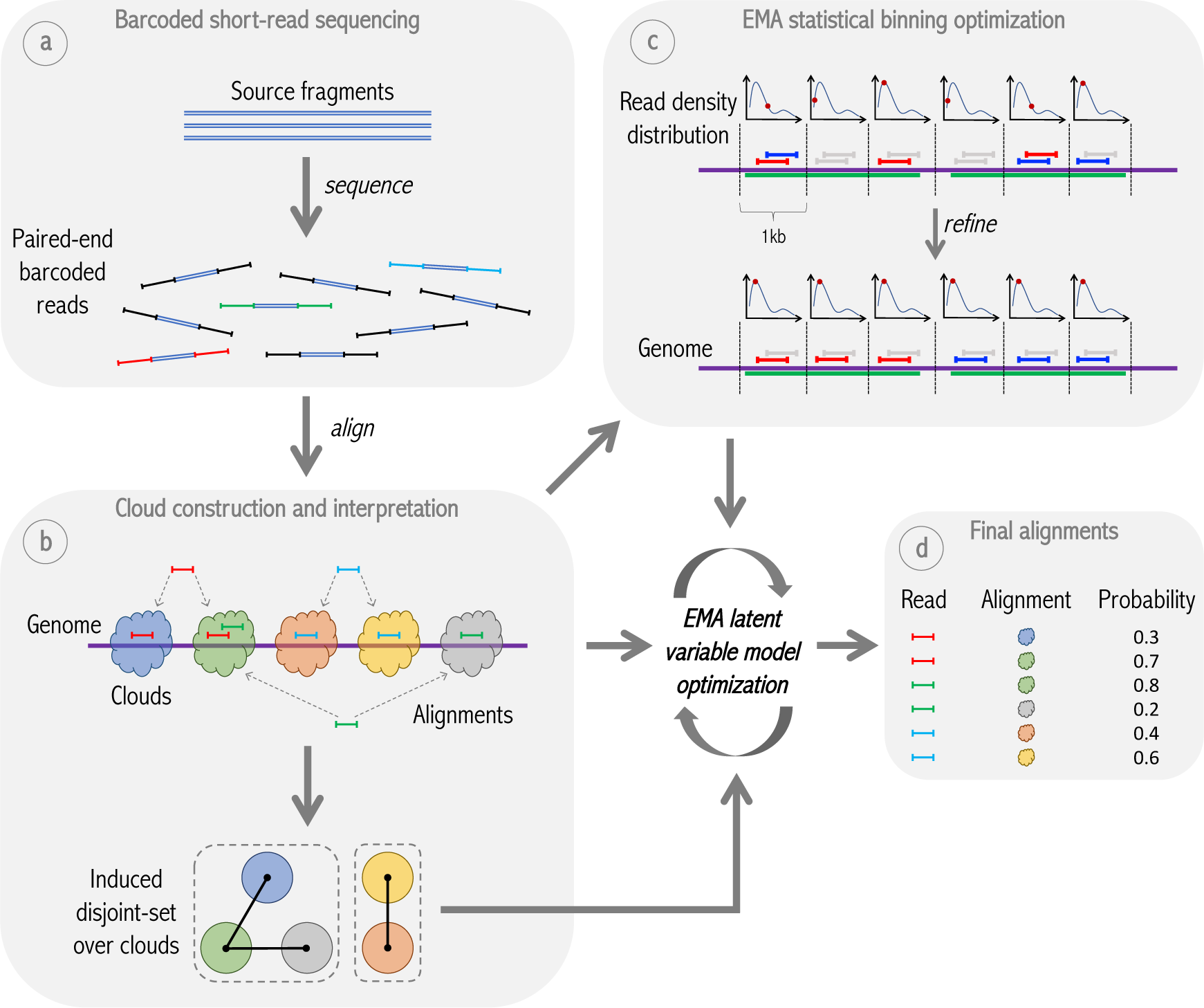
Overview of EMA pipeline. **(a)** Idealized model of barcoded short-read sequencing, wherein some number of unknown source fragments in a single droplet are sheared, barcoded and sequenced to produce barcoded reads. **(b)** EMA’s “read clouds” are constructed by grouping nearby-mapping reads sharing the same barcode; these clouds represent possible source fragments. EMA then partitions the clouds into a disjoint-set induced by the alignments, where two clouds are connected if there is a read aligning to both; connected components in this disjoint-set (enclosed by dashed boxes) correspond to alternate possibilities for the *same* unknown source fragment. EMA’s latent variable model optimization is subsequently applied to each of these connected components individually. **(c)** EMA applies a novel statistical binning optimization algorithm to clouds containing multiple alignments of the same read to pick out the most likely alignment, by optimizing a combination of alignment edit distances and read densities within the cloud. In the figure, the green regions of the genome are homologous, thereby resulting in multi-mappings within a single cloud. **(d)** While the statistical binning optimization operates within a single cloud, EMA’s latent variable model optimization determines the best alignment of a given read between different clouds, and produces not only the final alignment for each read, but also interpretable alignment *probabilities* (see Figure 2).

By thinking of clouds not as arbitrary clusters of reads, but rather as distributions, we are able to produce more accurate alignments, and to assign interpretable probabilities to our alignments, which greatly improves downstream analyses. We demonstrate EMA’s performance by evaluating downstream genotyping and phasing accuracy using real 10x data. We found that genotypes called using EMA’s alignments contained over 30% fewer false positives than those called using Lariat’s alignments, and contained fewer false negatives as well, when run on independent 10x datasets of NA12878 and NA24385 using NIST GIAB high-confidence variant calls as a gold standard. Overall, we found that roughly 20% of all reads in our two datasets had multiple suitable alignments and were therefore able to be targeted by EMA’s optimization algorithm. Furthermore, we demonstrate that EMA successfully resolves alignments in the pharmacogenomically important and highly homologous *CYP2D* region through its statistical binning optimization, and was able to detect novel variants therein that remain undetected when using Lariat or BWA.

In addition to achieving superior accuracy, the EMA pipeline is 50% faster than Lariat— which translates into days faster for typical 10x datasets— and does not add any memory overhead to the alignment process. Thus, we expect the algorithms introduced here to be a fundamental component of read cloud-based methods in the future.

## 2 Methods

General barcoded read sequencing begins with splitting the source DNA into long fragments (10– 200kb) where each such fragment is assigned some barcode (a short 16bp DNA sequence). These fragments are sheared and each sheared piece has the assigned barcode ligated to it, whereupon standard short-read sequencing is applied to the sheared pieces. Each fragment is shallowly sequenced (i.e. fragment coverage is typically less than 1× in read depth, so each fragment position is covered by at most one read); an overall high coverage is attained by sequencing many different fragments covering the same genomic region. As a result of this process, barcoded reads have the same low error rates as typical Illumina whole-genome sequencing reads. An idealization of this process is illustrated in Figure 1a.

### 2.1 Standard data preprocessing

The first stage in the alignment process is to preprocess the data, for which we largely follow the same practices used by 10x Genomics’ WGS software suite, Long Ranger [11]. The purpose of this preprocessing is to:

- extract the barcode from the read sequence,
- error-correct the barcode based on quality scores and a list of known barcode sequences,
- and group reads by barcode into “barcode buckets” to enable parallelism during alignment.

In summary, in the barcode extraction stage, we remove the 16bp barcode from the first mate of each read pair, and trim an additional 7bp to account for potential ligation artifacts resulting from the barcode ligation process during sequencing (the second mate shares the same barcode as the first mate). Subsequently, we compare each barcode to a list *B* of known barcodes to produce a per-barcode count, and compute a prior probability for each known barcode based on these counts. Note that this list is designed such that no two barcodes are Hamming-neighbors of one another. Now for each barcode *b* not appearing in *B*, we examine each of its Hamming-1 neighbors *b*′ and, if *b*′ appears in *B*, compute the probability that *b*′ was the true barcode based on its prior and the quality score of the changed base. Similarly, for each *b* appearing in *B*, we consider each Hamming-2 neighbor *b*′ and compute the probability that *b*′ was the true barcode in an analogous way. Lastly, we employ a probability cutoff on the barcodes, and thereby omit the barcodes of reads that do not meet this cutoff. Any read not carrying a barcode after this stage is aligned with a standard WGS mapper such as CORA [6] or BWA [7].

While in standard read alignment parallelism can be achieved at the read-level, for barcoded read alignment we can only achieve parallelism at the barcode-level. Therefore, the last preprocessing step is to group reads by barcode into some number of buckets. Each such bucket contains some range of barcodes from *B*, which are all grouped together within the bucket. This enables us to align the reads from each bucket in parallel, and to merge the outputs in a post-processing step.

### 2.2 Latent variable model for aligning barcoded reads to clouds

Here we employ a latent variable model for determining the optimal assignment of reads to their possible clouds. A “cloud” is defined to be a group of nearby alignments of reads with a common barcode, thereby representing a possible source fragment [10]. We consider all the reads for an individual barcode simultaneously, all-mapping and grouping them to produce a set of clouds for that barcode (Figure 1b). The clouds are deduced from the all-mappings by grouping any two alignments that are on the same chromosome and within 50kb of one another into the same cloud, which is the same approach employed by Lariat. While this heuristic works well in the majority of cases, it can evidently run into issues if, for example, a single read aligns multiple times to the same cloud. We address such cases below in Section 2.3, but assume in the subsequent analysis that clouds consist of at most one alignment of a given read. (Overall, we found that roughly 20% of all reads mapped to multiple clouds and were thus able to be optimized by EMA.)

As notation, we will denote by *c* the set of alignments contained in a given cloud. We restrict our analysis to a single set of clouds 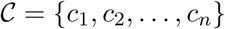 that corresponds to a connected component in the disjoint-set over clouds induced by alignments, as shown in Figure 1b (i.e. two clouds *c*_*i*_ and *c*_*j*_ will be connected if there is a read that has an alignment to both *c*_*i*_ and *c*_*j*_). Conceptually, the clouds in 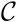 can be thought of as alternate possibilities for the *same* latent source fragment. By definition, for any given read aligning to some cloud in 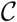, we will have to consider only the clouds in 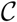 when determining the best alignment for that read, so we focus on each such set of clouds separately.

For 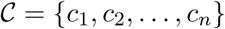, let *C*_*i*_ denote the event that cloud *c*_*i*_ represents the true source fragment. Since the clouds *c*_1_,…, *c*_*n*_ are different possibilities for the same source fragment, we have Pr(*C*_*i*_ ∩ *C*_*j*_) = 0 (*i* ≠ *j*) and 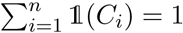 (where 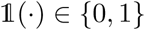 is an indicator for the specified event). We assume uniform priors on the clouds so that 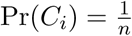 (while it is possible to devise a prior that takes into account features such as cloud length, we observed a large variance between clouds in our datasets that renders this unhelpful). Now, a cloud *c*_*i*_ can be conceptualized as an entity that generates some number of reads *K*_*i*_, parameterized by some weight 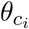, so that we can say 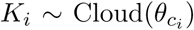 for some unknown “cloud” distribution over generated reads. We make the key assumption that, in expectation, 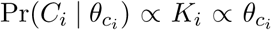 for all 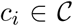. In other words, if a cloud is expected to have generated a large number of reads, then the probability that the cloud represents a true source fragment is high. Let 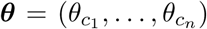 be the vector of cloud weights. We assume the cloud weights are normalized so that 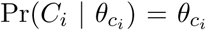, and that they are drawn from a uniform Dirichlet distribution so that ***θ*** ∼ Dir(**1**). Consider now the probability 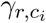 that a read *r* aligns to cloud *c*_*i*_ (denoted as an event by 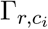) given the cloud parameters ***θ*** (i.e. 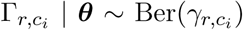), where Ber(*p*) is the Bernoulli distribution with parameter *p*). By Bayes’ rule, we can say:

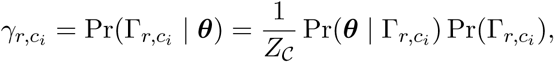

where 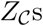 (and variants thereof) are normalization constants that are the same for each 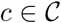. Since 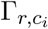 occurs if and only if *C*_*i*_ occurs, we have

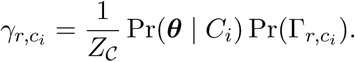

Applying Bayes’ rule again to Pr(***θ*** | *C*_*i*_) and using the fact that both Pr(***θ***) and Pr(*C*_*i*_) are uniform, we obtain

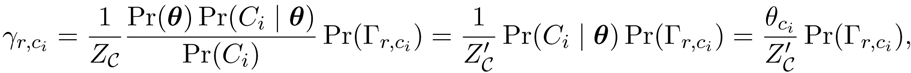

where 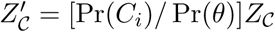. Note that 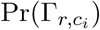 is a prior on the probability that *r* aligns to *c*_*i*_ that is not dependent on the barcode, but rather only on edit distance, mate alignment, and mapping quality as in standard short-read alignment. Henceforth, we refer to 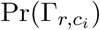 as 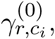, so that 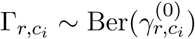.

Now we can form a prior 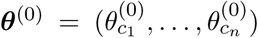 which is intuitively the initial vector of cloud weights. If we are given a set of alignment probabilities and a “current” ***θ*** estimate 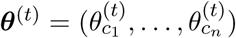 (initially *t* = 0), we can iteratively compute a better estimate ***θ***^(*t*+1)^ using the fact that 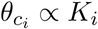 in expectation:

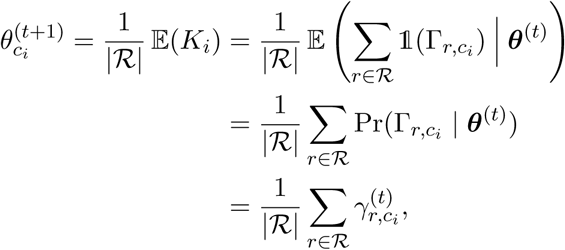

where ℛ is the set of reads mapping to any cloud in 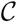, and the 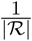 factor ensures that 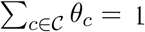. This latent variable model formulation naturally leads to an expectation-maximization algorithm— one of the two main ways of maximizing likelihood in such models— for determining the cloud weights and, thereby, the final alignment probabilities 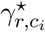. An implementation of this algorithm is given in the Appendix.

Each of the described variables and their interactions with one another is summarized in Figure 2. Once we determine the final alignment probabilities through this method (as in Figure 1d), we use them to compute mapping qualities (“MAPQs”), which are a standard per-alignment metric reported by all aligners and are frequently used by downstream analysis pipelines. Specifically, we take the MAPQ to be the minimum of the alignment probability, the barcode-oblivious alignment score and the MAPQ reported by BWA-MEM’s API (which is used in EMA’s current implementation to find candidate alignments). Importantly, we also report the actual alignment probabilities determined by EMA via a special standard-compliant SAM tag, so that they are available to downstream applications.

**Figure 2:**
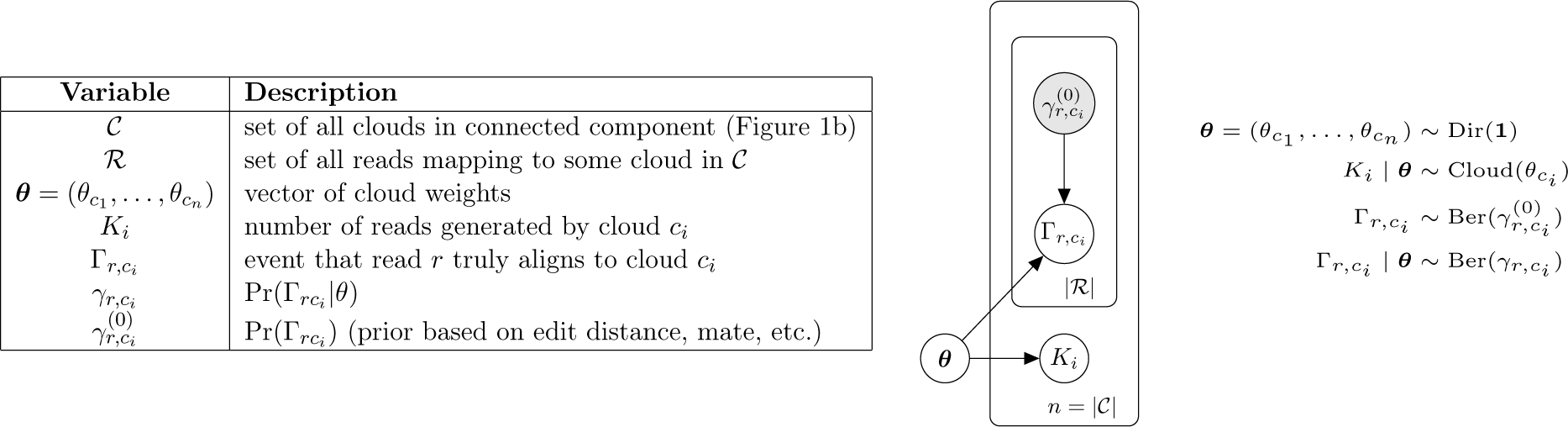
Graphical representation of EMA’s latent variable model involved in barcoded read alignment. ***θ*** denotes the vector of cloud weights; *K*_*i*_ denotes the number of reads generated by cloud 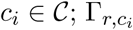 denotes whether read *r* ∈ ℛ maps to cloud *c*_*i*_, and 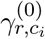 is a prior on this event based on barcode-oblivious information like edit distance, mate alignment, etc.

### 2.3 Statistical binning enables handling of multi-mappings in a single cloud

While the 50kb-heuristic described above is typically effective at determining the clouds, it does not take into account the fact that a single read may align multiple times to the same cloud (which can occur if a cloud spans two or more homologous regions). In such cases, rather than simply picking the alignment with lowest edit distance within the cloud, as is the current practice, we propose a novel alternative approach that takes into account not only edit distance but also read *density*. We take advantage of the insight that there is typically only a single read pair per 1kb bin in each cloud; the exact distribution of read counts per 1kb bin is shown in Figure 6 (Appendix). Now consider the case where one of our source fragments spans two highly similar (homologous) regions, and thereby produces a cloud with multi-mappings, as depicted in Figure 1c. If we pick alignments solely by edit distance, we may observe an improbable increase in read density (as shown in the figure). Consequently, we select alignments for the reads so as to minimize a combination of edit distance *and* abnormal density deviations.

Specifically, consider any cloud with multi-mappings consisting of a set of reads *R* = {*r*_1_,…, *r*_*n*_}, and denote by *A*_*r*_ the set of alignments for read *r* ∈ *R* in the cloud. Additionally, let *a*_*r*_ ∈ *A*_*r*_ denote the currently “selected” alignment for *r*. We will initially partition the cloud, spanning the region from its leftmost to its rightmost alignment, into the set of bins *B* = {*b*_1_,…, *b*_*n*_} of equal width *w*, where each bin *b*_*i*_ covers the alignments whose starting positions are located in the interval [*i* · *w*, (*i* + 1) · *w*), as shown in Figure 1c. In practice, we set *w* to 1kb. Denote by 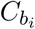 the random variable representing the number of reads in bin *b*_*i*_, where 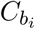 is drawn from the bin density distribution CloudBin(*i*). Lastly, let 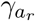 denote the prior probability that alignment *a*_*r*_ is the true alignment of the read *r* based on edit distance and mate alignments alone. Our goal is to maximize the objective:

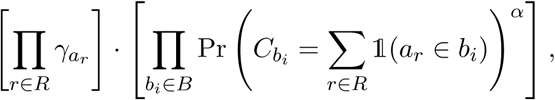

where *α* is a parameter that dictates the relative importance of the density probabilities compared to the alignment probabilities. We determine the distribution CloudBin(*i*) of each 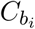 beforehand by examining uniquely-mapping clouds that we are confident represent the true source fragment. Taking the logarithm, this objective becomes:

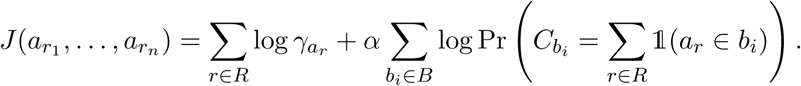

We optimize *J* through simulated annealing by repeatedly proposing random changes to *a*_*r*_ and accepting them probabilistically based on the change in our objective (the corresponding algorithm is described in the Appendix).

We apply the preceding latent variable optimization algorithm to deduce optimal alignments *between* clouds and, if necessary, use this statistical binning algorithm to find the best alignments *within* a given cloud.

## 3 Results

We compared the performance of EMA with Lariat [11] (10x’s own aligner and a component of the Long Ranger software suite) and BWA-MEM [7] (which does not take advantage of barcoded 10x data, and was therefore used as a baseline for what can be achieved with standard short-reads). In order to benchmark the quality of the aligners, we examined downstream genotyping accuracy, alignments in highly homologous regions, and downstream phasing accuracy.

We ran each tool on two 10x *H.sapiens* datasets for NA12878 and NA24385, and used the corresponding latest NIST GIAB [12, 13] high-confidence variant calls as a gold standard for each (version 3.3.2). For both EMA and BWA, we performed duplicate marking after alignment using Picard’s MarkDuplicates tool [14] (with barcode-aware mode enabled in the case of EMA); Long Ranger performs duplicate marking automatically. Genotypes were called by GATK’s HaplotypeCaller [15, 16] with default settings, while phasing was done by HapCUT2 [17] in barcode-aware mode. Genotyping accuracies were computed using RTG Tools [18].

Overall, we found that 20% of reads from our NA12878 dataset and 18% from our NA24385 dataset had multiple suitable alignments and were therefore able to be targeted by EMA’s optimization algorithm.

### 3.1 EMA improves downstream genotyping accuracy

Figure 3 shows genotyping accuracies for each aligner. For NA12878, EMA attains an *F*_1_ score of 0.944 compared to Lariat’s *F*_1_ of 0.925. For NA24385, EMA attains an *F*_1_ of 0.924 compared to Lariat’s 0.899. We found that for both datasets, EMA produced over 30% fewer false positive variant calls compared to Lariat, and produced fewer false negative calls as well. Interestingly, BWA-MEM (which does not take barcodes into account) performed marginally better than Lariat here, especially on the NA24385 dataset. Nevertheless, EMA also outperforms BWAMEM, attaining the fewest false positive and false negative variant calls between the three aligners on both datasets.

**Figure 3:**
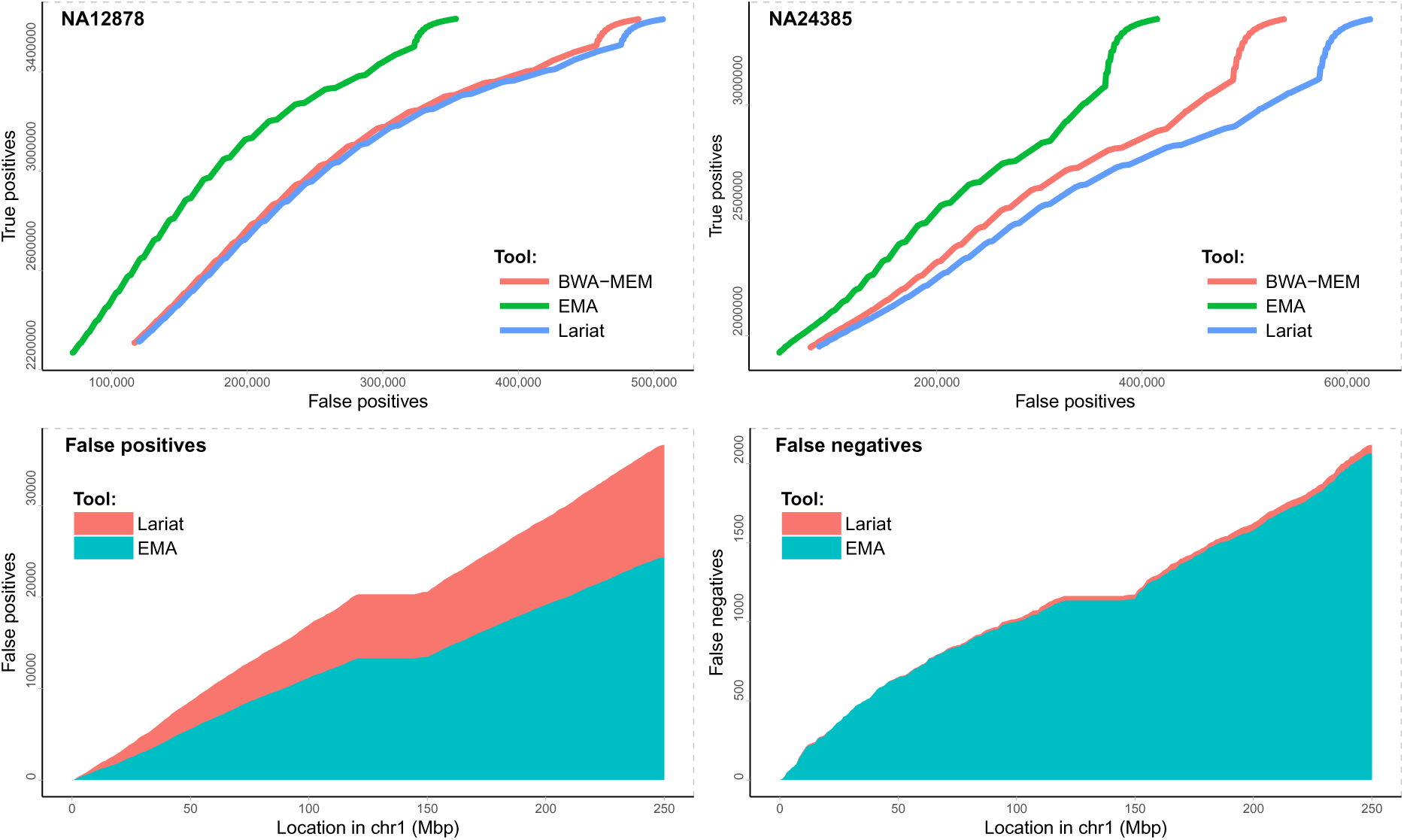
Genotyping accuracy for each aligner. The top row shows true positives as a function of false positives for alignments produced by EMA (green), Lariat (blue) and BWA-MEM (red) on the two well-studied samples NA12878 (left) and NA24385 (right). Genotype confidences are determined by the GQ (“genotype quality”) annotations generated by GATK’s HaplotypeCaller. The bottom row is a cumulative histogram of false positives (left) and false negatives (right) throughout chromosome 1 for NA12878. EMA achieves more than a 30% average improvement over the other methods.

### 3.2 EMA improves alignments in highly homologous regions

Among the principal promises of barcoded read sequencing is better structural variation detection, which invariably requires resolving alignments in homologous regions. One of the most important such regions is the *CYP2D* region in chromosome 22, which hosts *CYP2D6* — a gene of great pharmacogenomic importance [19]— and the two related and highly homologous regions *CYP2D7* and *CYP2D8*. The high homology between *CYP2D6* and *CYP2D7* makes copy number and variant calling in this region particularly challenging. The majority of aligners misalign reads in this region. This is especially evident in NA12878 which, in addition to the two copies of both *CYP2D6* and *CYP2D7*, contains an additional copy which is a fusion between these two genes [20], as well as *CYP2D7* mutations which introduce even higher homology with the corresponding *CYP2D6* region. Especially problematic is exon/intron 8 of *CYP2D6*, where many reads originating from *CYP2D7* end up mapping erroneously (see Figure 4 for a visualization). Even the naïve use of barcoded reads is not sufficient: both homologous regions in *CYP2D* are typically covered by a single cloud. For example, Lariat performs no better than BWA in this region (Figure 4). For these reasons, we chose to evaluate EMA in *CYP2D* to benchmark its accuracy in such highly homologous regions.

**Figure 4:**
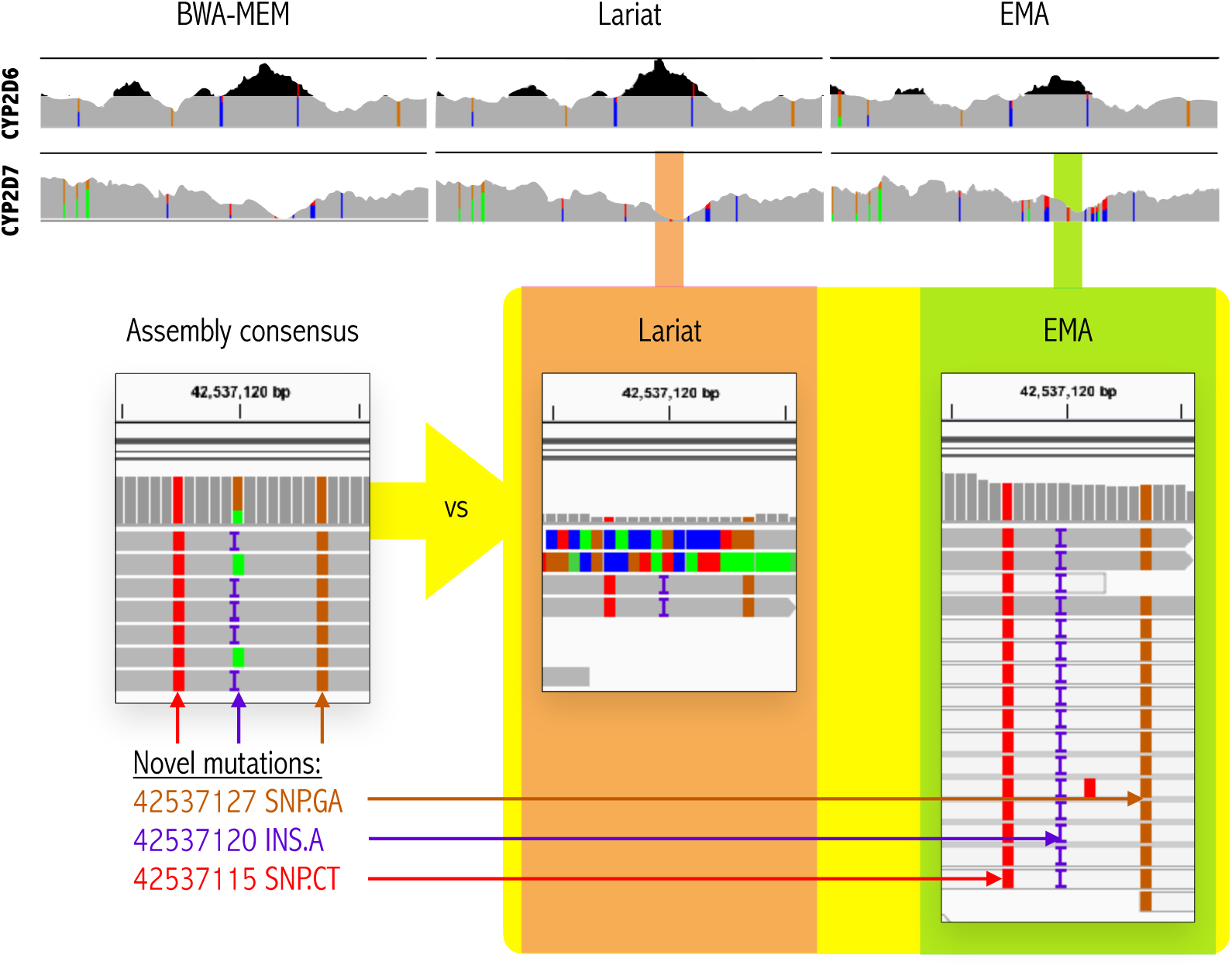
Positive effect of EMA’s statistical binning in the *CYP2D* region. The top image shows the read coverage for the region around exon/intron 8 within *CYP2D6* (top row) and *CYP2D7* (bottom row). Spurious coverage peaks (i.e. increases in observed coverage likely to be false) in *CYP2D6* are shaded black. EMA is clearly able to remove the problematic peaks and correctly assign them to *CYP2D7*. The bottom portion shows the newly assigned mappings to *CYP2D7*: EMA’s alignments agree with the assembly consensus sequence (observe the insertion and two neighboring SNPs detected by EMA). By contrast, both Lariat and BWA-MEM aligned virtually no reads to this region, and were thus unable to call these mutations.

As can be seen in Figure 4, EMA’s statistical binning strategy significantly smooths out the two problematic peaks in *CYP2D6* and *CYP2D7*. This technique enabled us to detect three novel mutations in *CYP2D7* (Figure 4) which exhibit high homology with the corresponding region in *CYP2D6*. Thus all reads originating from these loci get misaligned to *CYP2D6*, especially if one only considers edit distance during the alignment (as Lariat and BWA do). Such misalignments are evident in the “peaks” and “holes” shown in Figure 4. We cross-validated this region with the consensus sequence obtained from available NA12878 assemblies [21, 22, 23], and confirmed the presence of novel mutations.

As an aside, the copy number derived from EMA’s alignments in this problematic region (spanning from exon 7 to exon 9 in *CYP2D6* and *CYP2D7*) was closer to the “expected” copy number by 25% compared to the copy number derived from Lariat’s alignments (additional details are in the Appendix). Finally, statistical binning did not adversely impact phasing performance in this region, as we were able to correctly phase *CYP2D6*4A* alleles in our NA12878 sample from EMA’s alignments.

### 3.3 EMA improves downstream phasing

We applied HapCUT2 [17], which supports barcoded reads, to phase the variants called by GATK for both EMA’s and Lariat’s alignments. We evaluated our results with the phasing metrics defined in the HapCUT2 manuscript (Table 1). EMA provides more accurate phasings with respect to any metric in comparison to Lariat.

**Table 1:** Phasing results for EMA and Lariat on NA12878 and NA24385. Bold type indicates best results. Error metrics indicate the number of “incorrect” phasings compared to the GIAB gold standard; N50 metrics are based on the length of the phase blocks (bp).

| Sample | Tool | Switch errors | Mismatch errors | Flat errors | N50 |
| --- | --- | --- | --- | --- | --- |
| NA12878 | EMA | **12,796** | **14,163** | **538,169** | **111,392,359** |
|  | Lariat | 13,001 | 14,705 | 609,858 | 92,447,569 |
| NA24385 | EMA | **10,240** | **14,110** | **377,957** | **115,423,711** |
|  | Lariat | 10,472 | 14,655 | 429,896 | **115,423,711** |

### 3.4 EMA is computationally more efficient

Runtimes and memory usage for each aligner are given in Table 2 (for the NA12878 dataset). These times include alignment, duplicate marking and any other data post-processing (e.g. BAM sorting/merging). The reported memory usages are per each instance of the given mapper. In general, we found the full EMA pipeline to be about 1.5× faster than Lariat.

**Table 2:** Runtime and memory usage of each alignment tool on NA12878 dataset. Numbers in parenthesis indicate the performance of the aligner alone (i.e. without sorting, merging or duplicate marking). Each mapper was allocated 40 threads on an Intel Xeon E5-2650 CPUs @ 2.30GHz.

| Tool | Time (hh:mm) | Memory (GB) |
| --- | --- | --- |
| EMA | 14:58 (10:40) | 5.4 |
| Lariat | 21:49 (12:45) | 7.0 |
| BWA-MEM | 14:49 (9:52) | 5.5 |

## 4 Conclusion

EMA applies a latent variable model to the barcoded read alignment problem, and in so doing outperforms the state-of-the-art in virtually every metric. By thinking of clouds not as arbitrary clusters of reads, but rather as *distributions*, we are able to not only produce more accurate alignments, but also to assign interpretable probabilities to our alignments. This ability has several benefits, the most immediate of which is that it enables us to set a meaningful confidence threshold on alignments. Beyond this, these alignment probabilities can be incorporated into downstream applications like genotyping, phasing and SV detection. We demonstrate this here by computing mapping qualities based on these probabilities, which are then used in genotyping and phasing; yet specialized algorithms centered around these probabilities are also conceivable.

Moreover, EMA is able to effectively discern between multiple alignments of a read in a single cloud through its statistical binning optimization algorithm. This addresses one of the weaknesses of barcoded read sequencing as compared to long-read sequencing; namely, only a relatively small subset of the original source fragment is observed— and more specifically, that the order of reads within the fragment is not known— making it difficult to produce accurate alignments if the fragment spans homologous elements. By exploiting the insight that read densities within a fragment follow a particular distribution, EMA more effectively aligns the reads produced by such fragments, which can overlap regions of phenotypic or pharmacogenomic importance, like *CYP2D* as demonstrated above.

As we usher in the new wave of next-generation sequencing technologies, barcoded short-read sequencing will undoubtedly play a central role, and fast and accurate methods for aligning barcoded reads such as those presented here will ultimately prove invaluable.

## Acknowledgements

We thank Chris Whelan, Chad Nusbaum, Eric Banks, as well as the rest of the GATK SV Group from the Broad Institute for providing us with data samples and many valuable suggestions. Also, we thank Jian Peng for helpful discussions, as well as Lillian Zhang for her help evaluating EMA’s performance.

## A Appendix

### A.1 Algorithms

#### Algorithm 1

Barcoded read alignment via expectation-maximization

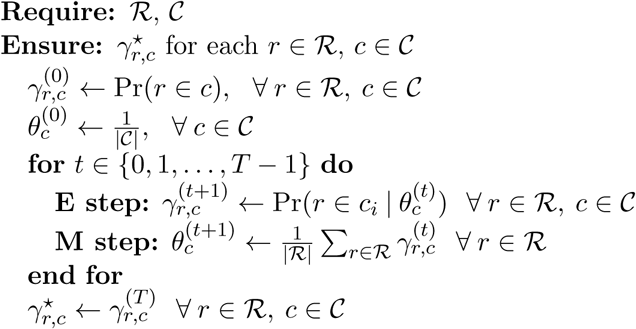

#### Algorithm 2

Read density optimization via simulated annealing

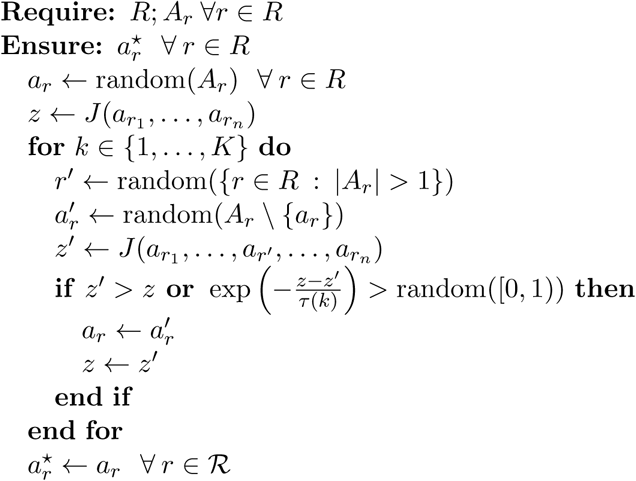

In Algorithm 2, *K* is the number of simulated annealing iterations, and *τ*(*·*) defines the annealing schedule (which can be taken to be an exponentially decreasing function). Other variables are described in detail in the main text.

### A.2 EMA implementation and parameters

A visualization of the EMA pipeline is given in Figure 5. The following parameters for EMA were used in the various experiments:

**Figure 5:**
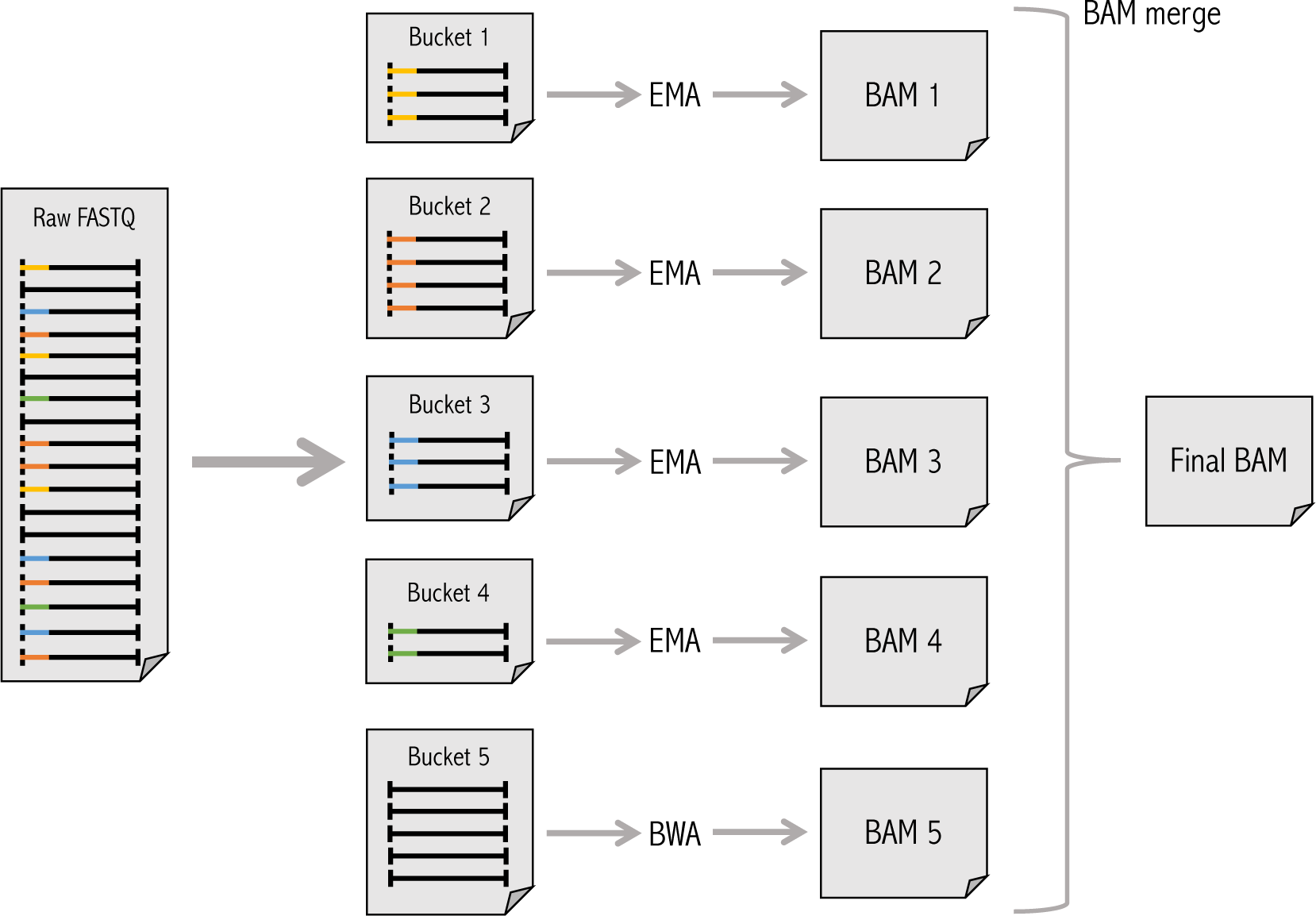
EMA pipeline. Raw FASTQs are split into buckets by barcode during preprocessing, then each bucket is processed by a separate instance of EMA in parallel. A special bucket containing non-barcoded reads is processed with BWA-MEM. The resulting BAM files are subsequently marked for duplicates and merged to produce a single, final BAM file as output.

| Parameter | Description | Value |
| --- | --- | --- |
| $T$ | number of EM iterations | 5 |
| $\alpha$ | density probability weight<br>in statistical binning | 0.05 |

EMA uses BWA-MEM’s C API to find candidate alignments just as Lariat does. EMA’s full code is available at http://ema.csail.mit.edu.

### A.3 Extended results

The following genotyping accuracy results were outputted by RTG Tools’ “vcfeval” utility after genotyping with GATK’s HaplotypeCaller (best results in each category are in bold):

| NA12878 |  |  |  |  |  |  |  |  |
| --- | --- | --- | --- | --- | --- | --- | --- | --- |
| Tool | True pos. baseline | True pos. call | False pos. | False neg. | Prec. | Sens. | $F_1$ | |
| EMA | **3,614,882** | **3,614,969** | **354,829** | **76,274** | **0.911** | **0.979** | **0.944** |  |
| Lariat | 3,613,361 | 3,613,447 | 507,666 | 77,795 | 0.877 | **0.979** | 0.925 |  |
| BWA-MEM | 3,613,352 | 3,613,443 | 489,605 | 77,804 | 0.881 | **0.979** | 0.927 |  |

| NA24385 |  |  |  |  |  |  |  |  |
| --- | --- | --- | --- | --- | --- | --- | --- | --- |
| Tool | True pos. baseline | True pos. call | False pos. | False neg. | Prec. | Sens. | $F_1$ | |
| EMA | **3,375,423** | **3,375,593** | **416,442** | **137,178** | **0.890** | **0.961** | **0.924** |  |
| Lariat | 3,374,059 | 3,374,236 | 624,103 | 138,542 | 0.844 | **0.961** | 0.899 |  |
| BWA-MEM | 3,374,670 | 3,374,845 | 539,915 | 137,931 | 0.862 | **0.961** | 0.909 |  |

### A.4 CloudBin distribution

The distribution of read counts in 1kb windows within the clouds is given in Figure 6.

**Figure 6:**
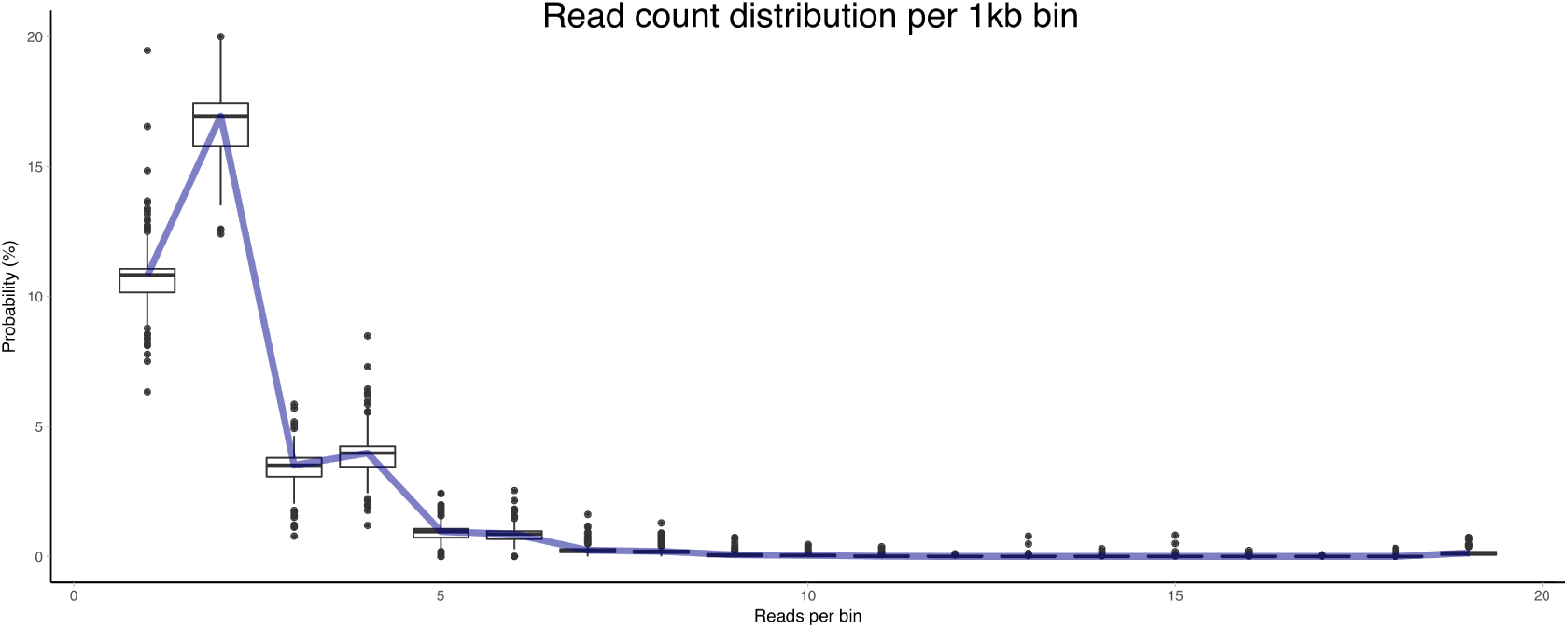
Distribution of the number of reads in a 1kb window within a cloud (based on NA12878 10x data). The box plots correspond to different bin offsets within the cloud.

### A.5 *CYP2D* analysis

The copy number of each intron and exon in the *CYP2D* region was obtained by running Aldy (http://cb.csail.mit.edu/cb/aldy/) on both Lariat’s and EMA’s alignments. We calculated the absolute difference from the estimated copy number for exon 7, intron 7, exon 8 and intron 8 (in both *CYP2D6* and *CYP2D7*), and the expected coverage (obtained from [20]: 2 for *CYP2D6* and 3 for *CYP2D7* regions). This difference is 7.51 for EMA’s alignments, and 10.22 for Lariat’s, implying an improvement of 25% if one uses EMA.

Furthermore, phased data from both Lariat’s and EMA’s alignments correctly linked *CYP2D6*4A* mutations together (i.e. chr22:42,524,947 C>T, chr22:42,525,811 T>C, chr22:42,525,821 G>T and chr22:42,526,694 G>A).

### A.6 Versions and parameters for other tools

| Tool | Version | Parameters |
| --- | --- | --- |
| Long Ranger | 2.1.6 | default |
| BWA | 0.7.15 | default |
| GATK HaplotypeCaller | 3.8.0 | default |
| HapCUT2 | eb3b64b | barcoded read mode |
| Picard MarkDuplicates | 2.9.2 | READ_ONE_BARCODE_TAG=BX<br>READ_TWO_BARCODE_TAG=BX |
| Samtools | 1.3.1 | n/a |
| RTG Tools | 3.8.4 | n/a |

